# Recurrent acquisition of nuclease-protease pairs in antiviral immunity

**DOI:** 10.1101/2025.07.28.667249

**Authors:** Owen T. Tuck, Jason J. Hu, Benjamin A. Adler, Claire E. O’Brien, Santiago C. Lopez, Kendall Hsieh, Charlotte Meredith, Erin E. Doherty, Arushi Lahiri, Jennifer A. Doudna

**Affiliations:** Department of Chemistry, University of California, Berkeley; Berkeley, USA; Innovative Genomics Institute, University of California, Berkeley; Berkeley, USA; Department of Molecular and Cell Biology, University of California, Berkeley, Berkeley, USA; California Institute for Quantitative Biosciences (QB3), University of California, Berkeley; Berkeley, USA; Howard Hughes Medical Institute, University of California, Berkeley; Berkeley, USA; Molecular Biophysics and Integrated Bioimaging Division, Lawrence Berkeley National Laboratory; Berkeley, USA; Li Ka Shing Center for Genomic Engineering, University of California, Berkeley; Berkeley, USA; Gladstone Institute of Data Science and Biotechnology; San Francisco, USA; Gladstone-UCSF Institute of Genomic Immunology; San Francisco, USA

## Abstract

Antiviral immune systems diversify by integrating new genes into existing pathways, creating new mechanisms of viral resistance. We identified genes encoding a predicted nuclease paired with a trypsin-like protease repeatedly acquired by multiple, otherwise unrelated antiviral immune systems in bacteria. Cell-based and biochemical assays revealed the nuclease is a proenzyme that cleaves DNA only after activation by its partner protease. Phylogenetic analysis showed that two distinct immune systems, Hachiman and AVAST, use the same mechanism of proteolytic activation despite their independent evolutionary origins. Examination of nuclease-protease inheritance patterns identified caspase-nuclease (*canu*) genomic loci that confer antiviral defense in a pathway reminiscent of eukaryotic caspase activation. These results uncover the coordinated activities of pronucleases and their activating proteases within different immune systems and show how coevolution enables defense system innovation.

## Introduction

Continual infection by viruses known as bacteriophage (phage) drives the evolution of widespread antiphage immune systems in bacteria and archaea^1–4^. Evolutionarily independent antiphage systems including CRISPR-Cas^5^, CBASS^6^, AVAST^2^, pAgo^7^, retrons^8^, Lamassu^4^ and Hachiman^9,10^ employ a small, interchangeable pool of genes that are typically fused to core immunity genes to provide new or alternative functionalities^3,11–14^. The acquisition of such fusion proteins represents a common but underappreciated mechanism of immune diversification through modular evolution^3,11–14^. However, it remains unclear whether newly acquired genes confer similar antiviral activities in the context of distinct antiviral systems, and whether gene acquisition might enable the discovery of new defense pathways^13,15^.

We identified multiple independent examples of bacterial immune systems with co-occurring genes encoding a trypsin-like protease and a predicted nuclease belonging to the metallo-β-lactamase (MBL)-type hydrolase superfamily ^16–18^. In bacterial antiphage systems, MBL hydrolases resemble nucleases from DNA uptake pathways^19,20^. While the predicted nuclease is always encoded by a separate, typically upstream gene, the protease occurs as a fusion with different antiviral defense-associated enzymes. First described in AVAST (antiviral ATPase/NTPase of the STAND superfamily, Avs)^2^, the nuclease-protease gene pair also occurs in Lamassu^4^, Hachiman and DRT (defense-associated reverse transcriptase) antiviral systems^21^. How encoded nuclease and protease enzymes contribute to defense in diverse antiviral contexts is unknown.

Here we present evidence for a unified nuclease-protease mechanism common to multiple antiviral systems with different infection-sensing mechanisms in bacteria. Cell-based and biochemical assays show that the nuclease exists initially as an enzymatically inactive ‘gated’ proenzyme. Site-specific cleavage by the partner protease liberates the nuclease active site. We show the activated nuclease is specific for DNA and executes indiscriminate DNA cleavage to restrict viral replication. This mechanism is similar in otherwise unrelated immune systems, suggesting that coordination between the nuclease and protease is conserved. Phylogenetic analysis revealed that gated nuclease homologs additionally exist in genomic loci alongside caspase-family proteases. We found that these caspase-nuclease (*canu)* loci encode defense systems in which the caspase activates the downstream nuclease, suggesting parallel coevolution of nucleases with multiple distinct partner proteases. Beyond revealing how paired nuclease and protease enzymes function in multiple antiviral systems, these results demonstrate that modular immune domains can guide the discovery and functional annotation of immune pathways, including systems like *canu* that display unexpected parallels with eukaryotic caspases.

## Results

### Repeated acquisition of paired genes in multiple antiphage systems

We noticed that some Hachiman, AVAST (antiviral ATPase/NTPase of the STAND superfamily, Avs), Lamassu and DRT (defense-associated reverse transcriptase) antiphage defense systems share a pair of co-occurring genes. In each case, a gene encoding a predicted nuclease (belonging to the metallo-β-lactamase (MBL)-fold hydrolase superfamily) sits immediately upstream of a gene encoding a trypsin-like protease fused to the N-terminus of a defense-associated enzyme (i.e. HamA, Avs1a or LmuA) (Fig. 1A). Across all four systems, the nuclease and protease co-occur and share sequence and structural homology (Fig. S1), suggesting they are evolutionarily and functionally linked.

**Fig. 1.**
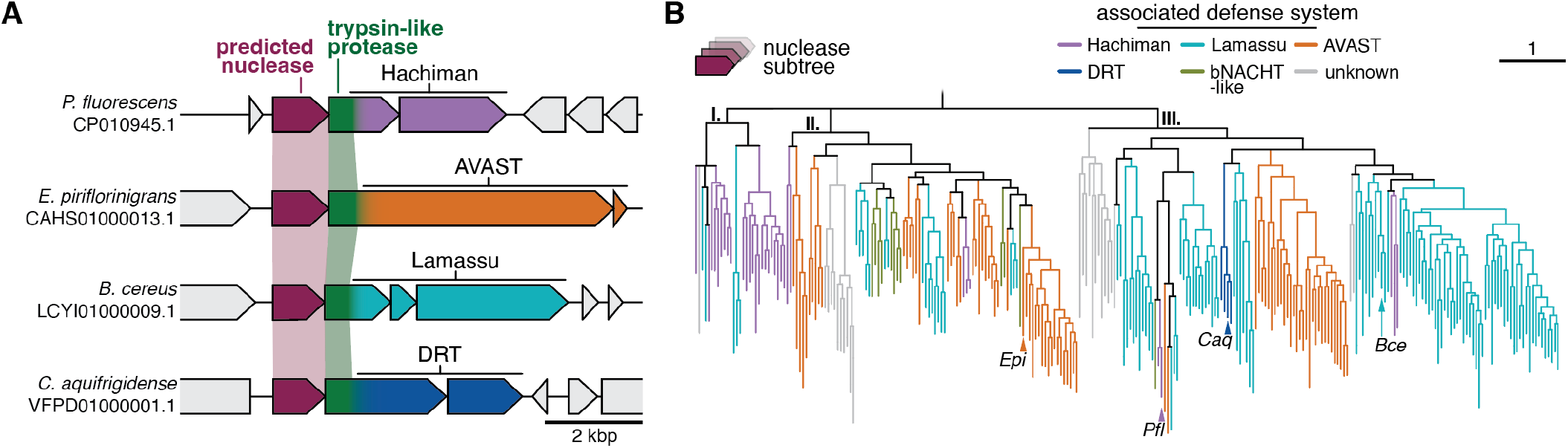
Multiple defense systems acquired a predicted nuclease and fused protease. **(A)** Loci illustrating the genetic architecture of multiple defense systems encoding a predicted N-terminal MBL hydrolase-like nuclease and a fused protease. **(B)** Phylogenetic tree of MBL hydrolase-like nucleases accessory to known defense systems which are associated with fused trypsin-like protease domain. Tree leaves are colored according to inclusion of the nuclease in known defense systems based on neighborhood analysis. Leaves shown in (A) are labelled. Branches with support values < 60 are deleted. A subtree is shown. For the complete phylogeny, see fig. S2A and B.

To determine how antiphage systems acquired nuclease-protease enzyme pairs, we performed sequence- and structure-based searches for homologs of the predicted nuclease, followed by genomic context mining (Fig. S2, see methods). Our analyses revealed a phylogenetic pattern consistent with frequent exaptation and horizontal genetic exchange. With the exception of rare DRT variants which emerged only once, all systems acquired the nuclease-protease pair on multiple independent occasions. For example, in clade III, a group of Hachiman-associated nucleases (HamM) diverged from a large radiation of Lamassu LmuA-H, and these sequences share a recent common ancestor with a similar cluster of nucleases from AVAST systems (Fig. 1B). We observed a similar branching order suggestive of frequent genetic exchange in clade II, where a group of HamM sequences resembles AVAST (Fig. 1B). Dissimilarity of the nuclease-protease pair across Hachiman loci provides additional evidence for their independent acquisition (Fig. 1B; Fig. S2C, S3). Our analyses reveal repeated transfer of paired nuclease and protease genes within known defense systems (Fig. 1B) and identify additional genomic loci with potential immune functions (Fig. S2C).

### Nuclease activation by a protease-Hachiman system

Our analyses indicate that prokaryotic immune systems frequently acquire nuclease-protease enzyme pairs from other, unrelated systems. To investigate how such acquisition alters immune system function, we began by examining examples of nuclease-protease gene integration into the Hachiman immune system. In ~5% of all Hachiman systems, the protease domain is present and fused to HamA, the nuclease responsible for activation-dependent DNA degradation (Fig. S4A). Analysis of diverse HamA sequences revealed multiple independent examples of protease-HamA fusions, consistent with the corresponding nuclease phylogeny (Fig. 1B; figs. S4B-G, S5A). Notably, other types of HamA fusions each occurred just once during Hachiman evolution, but do not co-occur with upstream nuclease genes and are predicted to confer diverse biochemical functions (Fig. S4 and Fig. S5A-C)^9,10,22^. Furthermore, in virtually all HamA fusions, the canonical D-GEXK catalytic motif required for HamA nuclease activity is absent (Fig. S5D). This evolutionary pattern implies that fusion Hachiman systems arose independently and lost HamA activity through mutation of active site residues. Domain fusion and nuclease loss are recurring themes in Hachiman evolution, and HamA is particularly predisposed to acquisition of the nuclease-protease pair.

We cloned and expressed the Pseudomonas fluorescens nuclease-protease Hachiman locus (*Pfl*HamMAB) in *E. coli* and verified that it provides defense against phage EdH4, reducing viral titer ~1000-fold (Fig. 2A, B; Fig. S6A). Single amino acid substitutions of predicted active site residues in the HamM predicted nuclease, the protease domain of HamA and the HamB helicase each abolished immunity, suggesting that nuclease, protease and helicase activities are all essential for antiphage activity. Despite toxicity upon expression, nuclease-protease Hachiman prevented cell lysis at low multiplicity of infection (MOI <0.1), while high viral doses (MOI >1) led to cell culture collapse (Fig. 2C). These findings suggest that *Pfl*HamMAB, like canonical HamAB systems, defends against phage by inducing programmed cell death or cell growth arrest.

**Fig. 2.**
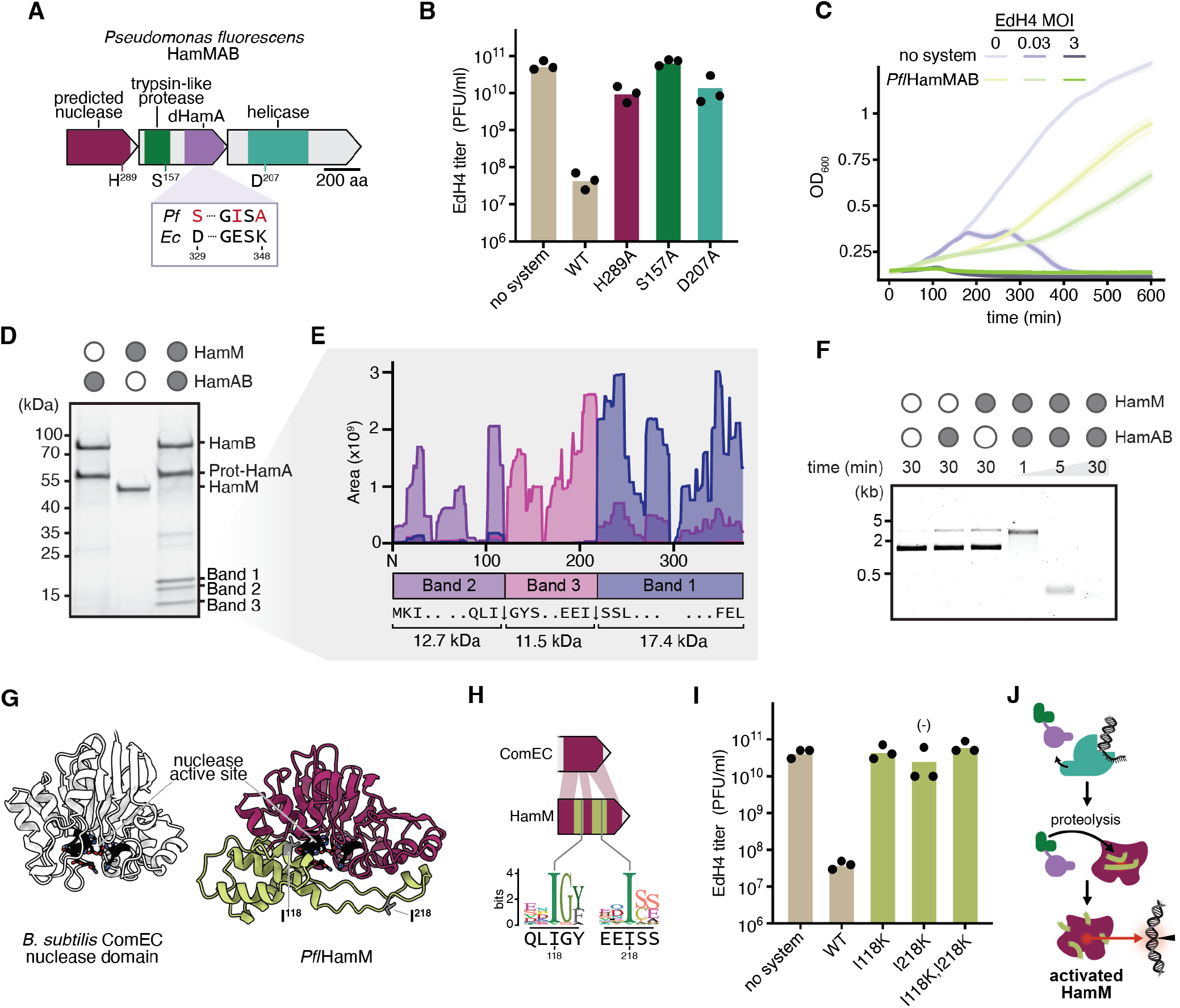
HamMAB encodes a specific protease which activates a partner nuclease. **(A)** *Pseudomonas fluorescens* HamMAB operon (CP010945.1) annotated with predicted catalytic residues and comparison to ECOR31 HamA active site. **(B)** Quantification of phage EdH4 plaque assays against *Pfl*HamMAB and mutants. Individual data points of three independent biological replicates are shown along with the mean. **(C)** Growth curves of *E. coli* expressing HamMAB and empty control during EdH4 infection at specified multiplicity of infection (MOI). Data are shown as mean ± standard deviation (shaded area) across three independent biological replicates. **(D)** In vitro reactions of HamM and HamAB visualized on a Coomassie PAGE gel. **(E)** Mass spectrometry reads of fragment 1, fragment 2, and fragment 3 mapped onto the HamM sequence. **(F)** In vitro plasmid clearance assay of HamM and HamAB with time course. **(G)** Comparison of AlphaFold3 models of the *Bacillus subtilis* ComEC MBL hydrolase-like nuclease domain (uniprot: P39695) with HamM, with active site colored black, protease cut sites colored gray, and insertions colored lime green. (H) Top, cartoon showing location of insertions in the HamM primary sequence and proteolysis sites. Bottom, sequence logos of cut site position among related nucleases, with the *Pfl*HamM primary sequence as a reference. **(I)** Quantification of phage EdH4 plaque assays against *Pfl*HamMAB and protease cut site mutants. Individual data points of three independent biological replicates are shown along with the mean. **(J)** Model for HamMAB defense.

To determine how nuclease-protease activities enable phage defense, we purified the HamM protein and the HamAB protein complex separately and tested their interactions in vitro (Fig. S7). Incubation of HamM with HamAB resulted in the appearance of three proteolytic fragments and the disappearance of a 42 kDa species corresponding to full length HamM (Fig. 2D). Mass spectrometry analysis mapped these fragments to specific regions of HamM, with two cleavage sites occurring immediately C-terminal to isoleucine residues I118 and I218 (Fig. 2E). Proteolysis was absent when we incubated HamAB with other proteins (Fig. S8). These data support the conclusion that the trypsin protease domain of HamA functions as an endopeptidase specific for certain isoleucine residues in HamM. Functional assays further demonstrated that plasmid incubation with either HamM or HamAB alone induced plasmid nicking, whereas incubation with HamM and HamAB together resulted in complete plasmid degradation (Fig. 2F). HamM together with HamAB does not degrade RNA (Fig. S9). These findings support the conclusion that proteolytic cleavage of HamM by the protease-HamA converts HamM into a potent deoxyribonuclease.

To understand how proteolysis activates HamM nuclease activity, we compared the AlphaFold predicted structure of HamM to a homologous MBL hydrolase-type nuclease involved in natural competence^19,23^. Superposition suggested that HamM includes two disordered insertions relative to the canonical nuclease, both of which appear to block the nuclease active site (Fig. 2G; Fig. S2). One cleavage-site isoleucine is located on each insertion; both are highly conserved at their respective positions (Fig. 2H). Hypothesizing that proteolysis removes the active site blockade created by insertions, we made single and double amino acid substitutions of cleavage site residues and tested phage defense. Mutation of either isoleucine residue is sufficient to abolish defense (Fig. 2I; Fig. S6B). Our results suggest that HamM is a ‘gated’ proenzyme that acquired regulatory insertions responsive to a partner protease. As HamB possesses ATPase activity against DNA substrates (Fig. S10), we propose that upon DNA sensing, HamB releases protease-HamA to trigger proteolytic cleavage of HamM and unleash its nuclease activity (Fig. 2J)^9,10^.

### Gated nuclease activation is conserved in AVAST

We wondered whether nuclease-protease enzyme partners retain functional properties despite their presence in divergent defense systems. To test this possibility, we investigated the nuclease-protease pair in an AVAST type 1 antiphage system from *Erwinia piriflorinigrans* (Fig. 1), an antiphage system with otherwise no homology to Hachiman and which was previously shown to provide phage defense in *E. coli*^2,24^. Structural prediction of the AVAST nuclease homolog (*Epi*Avs1a) shows dual insertions that appear to block the conserved active site, including two conserved isoleucine residues, analogous to structural features of *Pfl*HamM (Fig. 3A). We constructed single and double amino acid substitutions of conserved isoleucine residues and tested for AVAST-mediated defense against phage P1 in *E. coli*^2^. Each mutation alone reduced defense, while the double mutation eliminated AVAST-mediated protection entirely (Fig. 3B; Fig. S11). These data suggest that AVAST also activates a ‘gated’ effector nuclease via proteolysis, despite considerable phylogenetic distance from *Pfl*HamMAB (Fig. 1B). Conservation of this activation mechanism implies that the nuclease-protease pair behaves similarly in other antiphage systems (Fig. 3C). Consistent with this prediction, disordered loops proximal to the active site are present in every antiphage trypsin-associated nuclease identified in this study (Fig. S12).

**Fig. 3.**
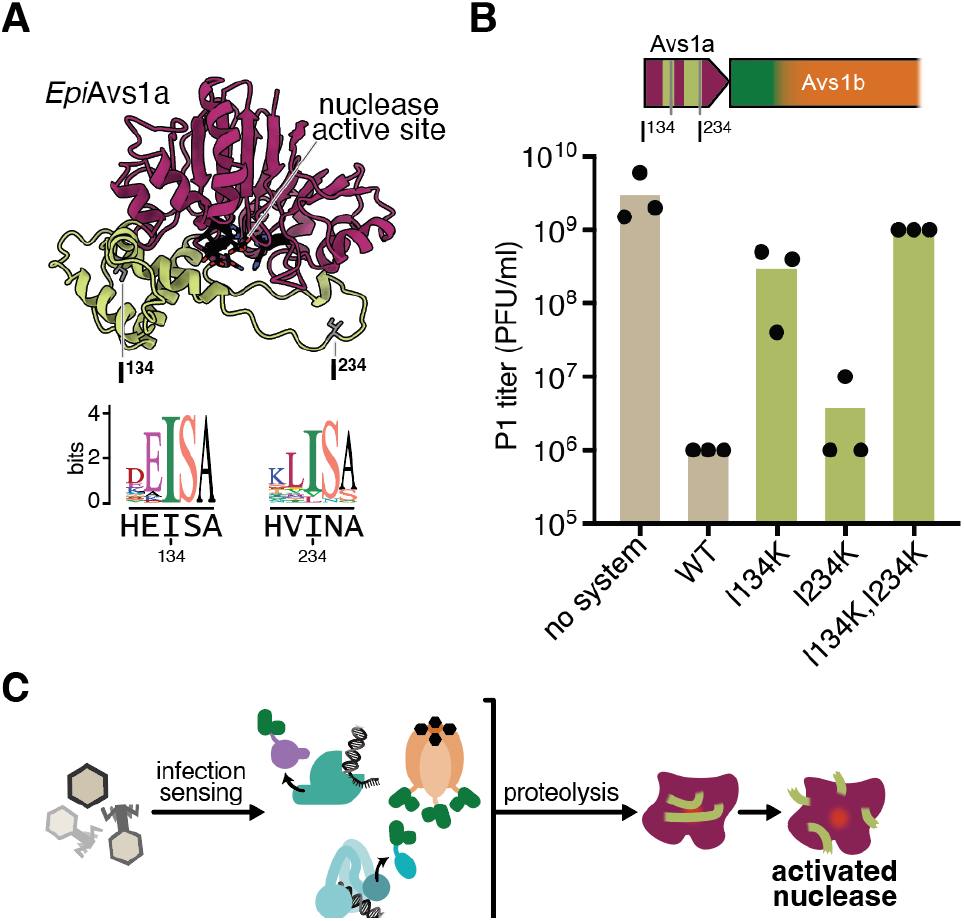
Nuclease-protease pairs function in AVAST. **(A)** AlphaFold3 model of the AVAST type 1 Avs1a nuclease from *E. piriflorinigrans* (CFBP5888), with active site colored black, predicted protease cut sites colored gray, and insertions colored lime green. Below, sequence logos of predicted cut site positions among related nucleases, with the *Epi*Avs1a primary sequence as a reference. **(B)** Quantification of phage P1 plaque assays against *Epi*AVAST and predicted protease cut site mutants. Individual data points of three independent biological replications are shown along with the mean. A portion of the locus showing locations of insertions in the *Epi*Avs1a primary sequence and predicted cut sites is shown. **(C)** Model for a conserved mechanism of pronuclease activation among known defense systems.

### Identification of a caspase-nuclease defense system

Gated nucleases co-occur with fused trypsin proteases in multiple known defense systems (Fig. 1A, B). Our phylogenetic analyses unexpectedly revealed an additional radiation of nuclease homologs that co-occur with proteases belonging to other diverse families (Fig. 4A; Fig. S2). The largest clade in this group includes a nuclease-encoding gene positioned downstream of a gene encoding a protease most closely related to caspases, which are essential to human innate immunity and apoptosis^18^. (Fig. 4A, B). These loci are variable in organization, encoding several genes of unknown function and multiple domains associated with oligomerization and protein-protein interactions, including tetratricopeptide repeats (TPR) and immunoglobulin-like (Ig-like) folds (Fig. 4B)^25^. Caspase-associated nucleases appear in immune hotspots known as ‘defense islands’ and near transposases, both signatures of immune function (Fig. S2C)^26^. We wondered whether these caspase-nuclease (*canu*) systems encode a similar mechanism, wherein the caspase activates its cognate nuclease to provide antiphage defense.

**Fig. 4.**
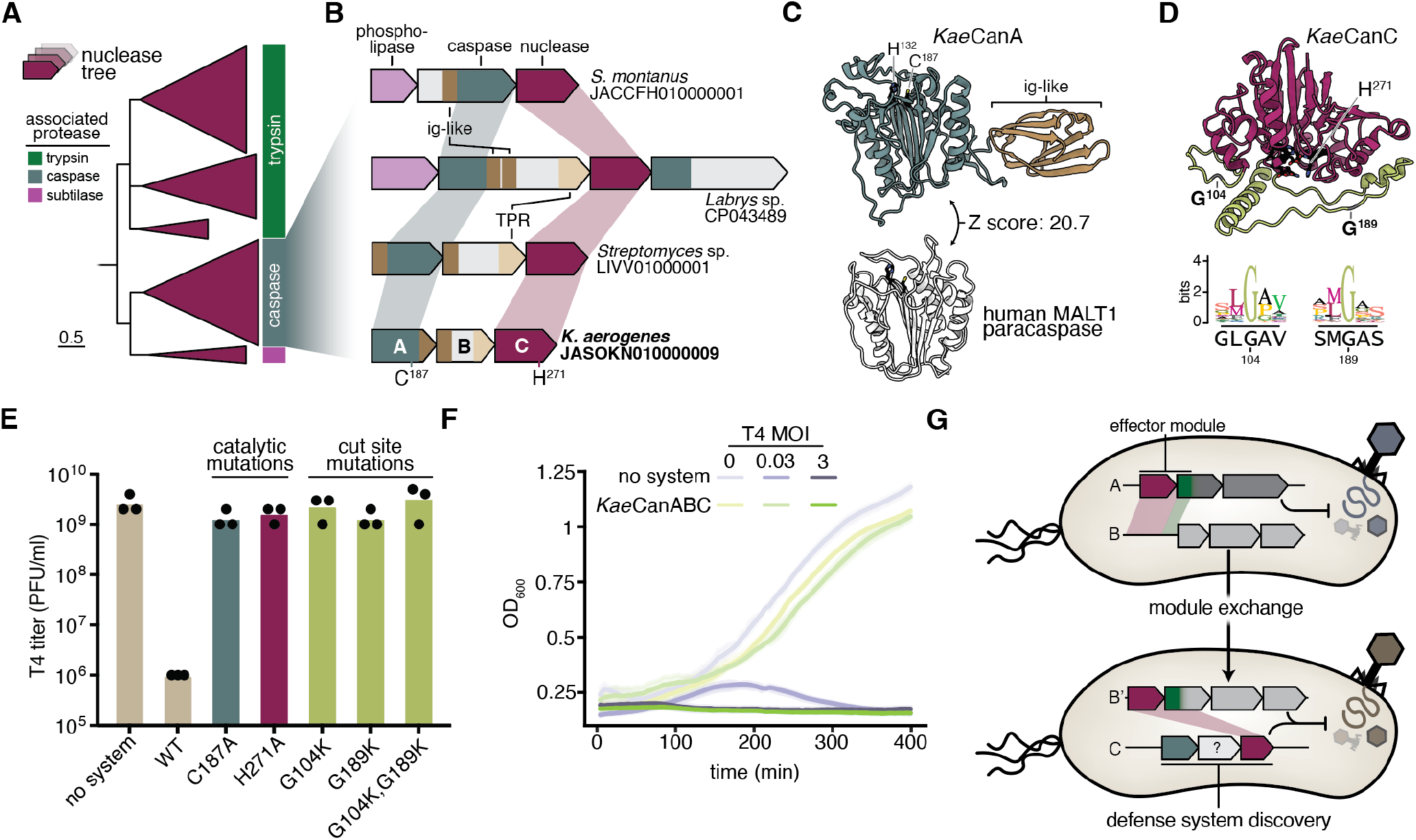
*Canu* is an antiphage defense system with a caspase-activated nuclease. **(A)** Phylogenetic tree showing major clades of nucleases associated with multiple protease families, as annotated by the outer track (right). A subtree is shown. For the complete phylogeny, see fig. S2. **(B)** Representative caspase-nuclease or *canu* loci with domains annotated (ig-like, immunoglobulin-like; TPR, tetratricopeptide repeat). The *Klebsiella aerogenes* CanABC operon (bottom) is annotated with predicted catalytic residues. **(C)** Structural homology (DALI Z-score: 20.7) between the AlphaFold3 model of *Kae*CanA and the human paracaspase MALT1 (Mucosa-associated lymphoid tissue lymphoma translocation protein 1, PDB: 3UOA), with active sites colored black. **(D)** AlphaFold3 model of *Kae*CanC, with active site colored black, predicted protease cut sites colored gray, and insertions colored lime green. Below, sequence logos of predicted cut site positions among related nucleases, with the *Kae*CanC primary sequence below. **(E)** Quantification of phage T4 plaque assays against *Kae*CanABC and mutants shown in (A). Individual data points of three independent biological replicates are shown along with the mean. **(F)** Growth curves of *E. coli* expressing *Kae*CanABC and an empty vector control during T4 infection at the specified MOI. Data are shown as mean ± standard deviation (shaded area) across three independent biological replicates. **(G)** Model for the exchange of the nuclease-protease pair (top) and a strategy to use coevolution and transfer to discover new defense loci (bottom).

In the prototypical three-gene *canu* system from *Klebsiella aerogenes* (*Kae*CanABC), the *canA* gene encodes a caspase domain structurally similar to the human MALT1 paracaspase, retaining the C-terminal Ig-like fold crucial for immune activation (Fig. 4B, C) (*27, 28*). The CanC nuclease active site, like *Pfl*HamM, *Epi*Avs1a and other nucleases in known defense systems, is likely blocked by dual disordered insertions based on structure prediction and comparison to related enzymes (Fig. 4D; fig S12). Both gating insertions contain conserved glycine residues (Fig. 4C), leading us to hypothesize that these positions are specifically cleaved by the caspase to activate nuclease activity.

To test *canu* activity, we challenged *E. coli* heterologously expressing *Kae*CanABC. We found that *Kae*CanABC provides strong protection against phage T4 (Fig. 4E; Fig. S13). Single amino acid substitutions of predicted active site residues in the caspase or nuclease domains abolished defense, suggesting that both catalytic activities are required. Mutation of either predicted proteolysis site in the nuclease was also sufficient to ablate defense. The *Kae*CanABC system overcame infection at low viral doses, but succumbed to phage when viral titer surpassed that of the host (Fig. 4F). Together, these data imply that *Kae*CanABC carries a conserved pronuclease which, when proteolytically processed by its partner caspase, protects cells from infection by programmed cell death or growth arrest.

## Discussion

In this study, we discovered that multiple different prokaryotic antiviral pathways employ related nucleaseprotease enzyme pairs to diversify immune function. Cellbased and biochemical data showed that proteasecatalyzed cleavage of a proenzyme form of the nuclease converts it to a DNase. Once activated, the nuclease destroys cellular DNA, likely killing or arresting infected cells to halt viral replication. Unlike most cell death effectors that act individually^2,6,29–31^, nuclease and protease genes function in tandem and co-evolve. This enzymatic partnership provides regulation that may safeguard against autoimmunity. For example, independent control over nuclease expression enables fine-tuning of its effect on cells, while fusion of its partner protease to core defense genes may prevent nonspecific activation^6,32–34^. Our results reveal that proteolytic activation, long recognized as central to eukaryotic immunity^35,36^, also shapes prokaryotic defense systems more extensively than previously appreciated^37–39^. Our phylogenetic analyses suggest that ancestral MBL hydrolase-like nucleases acquired autoinhibitory insertions containing endopeptidase sites, creating a requirement for protease partners to restore activity. The resulting nuclease-protease pairs were then transferred extensively between otherwise unrelated antiviral systems. Despite frequent mobilization, these enzyme pairs retain the coupled mechanism of nuclease activation, reflecting their common ancestry. Distinct genetic arrangements, including relative position and standalone versus fused configurations in trypsin and caspase systems, respectively, likely trace back to this fundamental functional association. Based on these observations, it is possible that other proteolytically activated enzyme classes participate in antiviral immunity, providing similar regulatory safeguards. Hallmarks of functional coupling, including protease coevolution and active site-blocking insertions, can guide the discovery of additional enzymatic partnerships.

One example of such a discovery is the identification of caspase-nuclease (*canu*) antiphage systems, which extend the diversity of known caspase-containing immune systems in prokaryotes^25,40–42^. In humans, multiple caspase paralogs orchestrate programmed cell death by either proteolytically activating other factors (initiator caspases), or by directly targeting cellular proteins (inflammatory and executioner caspases)^35,43,44^. Prokaryotes encode other caspase homologs with known functions in antiphage defense^40,42^, including Type III CRISPR^37–39^, gasdermin/CARD^45,46^, NLR^47^ and Thoeris systems^48^. *Canu* systems differ by encoding Ig-like domains that also occur in human caspases^25,27^. The *canu* system from *K. aerogenes* (*Kae*CanABC) provides potent phage defense and shows striking conservation of the nuclease activation mechanism found in Hachiman and AVAST. These results suggest that the *canu* caspase functions as a specific endopeptidase, analogous to eukaryotic initiator caspases^49^. While *canu* system diversity and functions await further exploration, conservation of nuclease activation provides a basis for determining the mechanism of *canu* defense.

## Supporting information

Supplementary References and Figures

## Acknowledgements

We thank Dr. T. Iavarone of the QB3/Chemistry Mass Spectrometry Facility, which received support from the NIH (1S10OD020062-01), and Dr. R. Maxwell of the Vincent J. Coates Proteomics/Mass Spectrometry Laboratory (RRID: SCR_025852) for assistance with mass spectrometry data acquisition. K. Zhou, J. Ye and E. Ma provided valuable technical advice and support. We thank K. Lucas for work in project oversight, facilitating collaboration, and supporting the scientific mission of the lab. We are grateful to K. Wasko, S. Swartz, P. Yoon and A. Guitor for critical feedback on this manuscript, and to all members of the Doudna Lab for helpful discussions. The Doudna Lab also receives support from NIH/NIAID (U54AI170792, UH3AI150552 and U01AI142817), NIH/NINDS (U19NS132303), NIH/NHLBI (R21HL173710), NSF (2334028), Lawrence Livermore National Laboratory, Apple Tree Partners (24180), UCB-Hampton University Summer Program, Mr. Li Ka Shing, Koret-Berkeley-TAU, Emerson Collective and the Innovative Genomics Institute (IGI).

## Author contributions

Conceptualization: OTT, JJH, BAA, JAD; Methodology: OTT, JJH, BAA, CEO, SCL, EED; Investigation: OTT, JJH, BAA, CEO, SCL, KH, CM, AL; Visualization: OTT, JJH, CEO, CM; Writing - original draft: OTT, JJH, JAD; Writing - revision, review & editing: OTT, JJH, BAA, CEO, SCL, KH, CM, EED, AL, JAD

## Materials and Methods

### Nuclease phylogenetic analysis

Phylogenetic analyses of MBL hydrolase-like nucleases were initiated by generating a seed alignment of HamM, Avs1a and LmuA genes using ClustalOmega^50^. The resulting alignment was manually trimmed and then used to generate a custom HMM with hmmbuild^51^, and subsequently hmmsearch was performed on the uniprot database (2025_01). Unique HMM hits with an e-value less than 1E-3 were selected, and clustered with MMseqs2 easy-linclust using --min-seq-id 0.9^52^. Multiple sequence alignment using ClustalOmega was performed, followed by manual trimming. Finally, IQ-TREE was performed with the following parameters: -bb 1000 -nm 1500 -safe, with ModelFinder used to select VT+F+R10^53,54^. The phylogenetic tree was visualized with iTOL^55^, with branches with bootstrap values below 60 deleted. A large clade containing the ComEC hydrolase/nuclease domain was chosen as a root. A subtree of only trypsin-like protease clades is shown in Fig. 1A, and a cartoon subtree of all protease-associated domains is shown in Fig 4A.

Context analysis was performed by extracting 10 kbp up- and downstream of hmmsearch hits using the NCBI Entrez tool^56^. PADLOC was then used to annotate defense systems within the gene neighborhood^57^. If hit proteins were encoded within a known defense system as determined by PADLOC, these were annotated as such in phylogenetic trees using iTOL annotation tools^55^. Leaves without defense system annotations were manually inspected and labelled. Nuclease structures from *Pfl*HamM, *Epi*Avs1a, *Bce*LmuA-H, and UG9 DRT systems were predicted using the AlphaFold3 web server^23^. Structures were aligned and visualized with matchmaker in ChimeraX^58^.

### Hachiman HamA phylogenetic analysis

For analysis of Hachiman diversity, unique proteins from InterPro protein family IPR014976 and annotated as Hachiman antiphage defense system protein HamA in Refseq and Genbank were pooled and chosen for phylogenetic analysis. MMseqs2 easy-linclust was run on pooled sequences with --min-seq-id 0.5^52^. Then, an HMM was built based on the clustered sequences with hmmbuild, and subsequently hmmsearch was performed on the genbank nr nucleotide database (v274)^51^. Unique HMM hits with an evalue less than 1E-5 were selected, and clustered with MMseqs2 easy-linclust using --min-seq-id 0.9. Multiple sequence alignment using muscle5.1 -super5 was performed on the clustered sequences, with an additional 10 homologs of the P. aquatile type IIS restriction endonuclease domain chosen as the outgroup^9,59^. Truncated Hachiman variants (HamA that are <150 amino acids) were manually removed and HamA-HamB fusions (likely due to stop codon misannotation) were removed from the sequence alignment, then re-aligned with muscle5.1 -super5. Poorly aligned regions were trimmed by trimAl with -gt0.2^60^. Finally, IQ-TREE was performed with --bb 1000 -m LG+F+G4^50^. The phylogenetic tree was visualized with iTOL^55^, with branches with bootstrap values below 60 deleted and branches annotated for taxonomy and the presence or absence of a fused domain (which were trimmed from the alignment to observe only HamA domain dynamics).

HamA were plotted by length, then uncommonly long HamA were extracted and aligned. Representatives of each long HamA sequence were folded using Colabfold^61^, then fused domains were manually extracted in ChimeraX^58^. Each of the five extracted N-terminal domains were used to search for structural homologs via the Dali search server^62^. Superimpositions were generated using the matchmaker function in ChimeraX^58^. Sequence logos and locus visualizations were generated using Geneious Prime.

### Bacterial strains and phages

For bacterial cultivation, *E. coli* strains maintained as 25% (v/v) glycerol stocks and stored at −80ºC were generally grown in LB Lennox media at 37ºC and 250 rpm aeration. Media was supplemented with carbenicillin (Sigma, 100 µg mL-1) or chloramphenicol (Sigma, 20 μg mL-1) where applicable. *E. coli* Top10 (invitrogen) was used as a cloning strain. For phage experiments using the *Pfl*HamMAB system, *E. coli* MDS42 was used. For *Epi*Avs and *Kae*CanABC phage experiments, we used *E. coli* K-12 MG1655. To purify proteins, *E. coli* BL21-AI (Invitrogen) was used.

Phages were propagated at 37ºC in LB Lennox media using an initial MOI of approximately 0.1 and host *E. coli* BW25113. After culture collapse, a drop of chloroform was added and cultures were centrifuged at 4000 g for 15 min. The supernatant was passed through a 0.2 µm filter and stored at 4ºC. All phage titers were determined on assay hosts with an empty vector control in the p15a backbone using plaque assays (described below).

### Plasmid construction

All plasmids used here were constructed using Gibson or Golden Gate assembly using PCR products purified by gel extraction (Zymo D4001). *Pseudomonas fluorenscens* and *Klebsiella aerogenes* loci were synthesized by Twist. For phage assays, all defense loci except AVAST were cloned under control of the pTet promoter in a p15a backbone encoding for chloramphenicol resistance The AVAST system, cloned from the source plasmid^2^, is under control of the native promoter in a p15a backbone, which encodes chloramphenicol resistance. For protein purification purposes, loci were cloned under a T7 lactose-controlled promoter in a high copy vector encoding carbenicillin resistance. All plasmids were sequence verified using Primordium. For a complete list of cloned sequences used in this study, see Table S1. A summary of plasmids used in this study and related sequences can be found in Table S2.

### Plaque assays

Plaque assays were performed as previously described, using the double agar overlay method^9^. In brief, 100 µL of saturated overnight cultures were added to molten LB Lennox agar (0.7% w/v agar, 60°C), then the mixture was supplemented with antibiotics and, in the case of *Pfl*HamMAB experiments, 2 nM anhydrotetracycline (aTc, Sigma) inducer. This mixture was poured over 1.5% w/v LB Lennox agar plates (presupplemented with antibiotic) and was allowed to cool and solidify. Phages were diluted tenfold in SM buffer (Teknova), then 2 µl of eight dilutions were spotted onto the top agar layer. After drying under sterile conditions, plates were incubated at 30ºC for 12-16 hours. Plates were then scanned and plaque forming units (PFU) were enumerated. Plaque assays were performed in biological triplicate, and data were analyzed using Graphpad Prism and Python.

### Liquid infection assays

To assay bacterial growth in the presence or absence of phage, OD600 was monitored in a 96-well plate format in a plate reader (Biotek Cytation 5). To initiate growth, saturated overnight cultures seeded in a microplate (Corning 3903) at a CFU of ~8e6 CFU per well in 200 μL of LB media. Phage were added to achieve the indicated MOI after separate dilution in SM buffer. Growth was monitored by measurement of OD600 every 5 minutes while shaking at 800 rpm (double orbital) at 30ºC for the indicated time span. Liquid growth assays were performed in biological triplicate, and data were analyzed using Python.

### Protein expression and purification

For complex purification vectors in the native locus format, only HamA was tagged with 10xHis-MBP-TEV. After transformation into BL21-AI *E. coli*, single colonies were inoculated in TB media and were grown to an optical density of ~0.6 then induced overnight at 16°C with 0.5 mM isopropyl-β-D-thiogalactopyranoside (IPTG) and 0.1% L-arabinose. Cells were harvested and resuspended in lysis buffer (20 mM 4-(2-hydroxyethyl)-1-piperazineethanesulfonic acid (HEPES) pH 7.7, 500 mM NaCl, 20 mM imidazole, 0.1% Triton X-100, 1 mM Tris (2-carboxyethyl)phosphine (TCEP), Complete EDTA (ethylenediaminetetraacetic acid)-free protease inhibitor (Roche), 0.5 mM phenylmethylsulfonyl fluoride (PMSF) and 10% glycerol). Cells were lysed by sonication, then clarified by centrifugation. The clarified lysate was incubated with Ni-NTA (Qiagen) resin for 1 hr. The resin was washed with wash buffer (20 mM HEPES, pH 8, KCl mM NaCl, 20 mM imidazole, 1 mM TCEP, and 5% glycerol), then bound protein was eluted with wash buffer supplemented with 300mM imidazole. Eluate was then run over an MBPTrap column (GE Healthcare), washed with MBP/SEC wash buffer (20 mM HEPES, pH 8, 150 mM KCl, 1 mM TCEP, and 5% glycerol), and eluted with MBP/SEC buffer supplemented with 10 mM maltose. Eluted protein from the MBPTrap column was treated with TEV protease overnight. Then, TEV protease-treated samples were run through Ni-NTA resin, and flow-through was collected. Flow-through was concentrated and run on a Superdex 200 10/300 GL column (Cytiva). Aliquots were concentrated and snap-frozen in liquid nitrogen for later use.

### Protease-HamAB proteolytic cleavage assay

Protease cleavage assays of HamM by protease-HamAB were performed in isothermal amplification buffer (20 mM Tris-HCl, pH 8.8, 10 mM (NH)_2_SO_4_, 50 mM KCl, 2 mM MgSO_4_, 0.1% Tween 20). 1.4 µM of protease-HamAB was incubated with 1 µM of HamM for 90 minutes at 37 ºC. Controls were performed with protease-HamAB only and HamM only as well. Reactions were terminated with addition of 4x Laemmli buffer. Samples were boiled for 3 minutes at 95ºC, loaded on a 4-20% SDS PAGE gel, stained by Coomassie InstantBlue, and imaged on a ChemiDoc MP (BioRad).

Protease cleavage assays of SSB (Fisher Scientific), RecBCD (New England Biolabs), and Topoisomerase I (New England Biolabs) by protease-HamAB were performed in isothermal amplification buffer. All reactions were performed with a final concentration of 0.25 μg/µL SSB, 0.5 units/µL RecBCD, and 0.25 units/µL Topoisomerase I. For the indicated reactions (Fig. S8), SSB, RecBCD, and Topoisomerase I were boiled at 95°C for 3 minutes before addition of protease-HamAB. Reactions were started by the addition of protease-HamAB to a final concentration of 0.5 µM and incubated at 30°C for 30 minutes. Controls with SSB, RecBCD, and Topoisomerase I only were conducted as well. Reactions were stopped by addition of 4x Laemmli buffer. Samples were boiled for 3 minutes at 95°C, loaded on a 4-20% SDS PAGE gel, stained by Coomassie InstantBlue, and imaged on a ChemiDoc MP (BioRad).

### In-gel mass spectrometry

Experiments for mass spectrometry were performed in MBP/SEC buffer supplemented with 1mM MgCl_2_. 9.6uM of protease-HamAB was incubated with 19.3uM of HamM for 60 minutes at 30°C. Reactions were terminated with addition of 4x Laemmli buffer. Samples were heated for 3 minutes, loaded on a 4-20% SDS PAGE gel, and visualized by Coomassie InstantBlue. Protein bands corresponding to proteolytic products of HamM were excised with a razor into 1 mm^2^ cubes. Gel pieces were washed twice with 50% (v/v) acetonitrile and 50mM ammonium bicarbonate (pH 8) for 15 minutes with shaking. Gel pieces were then dehydrated with 100% acetonitrile for 5 minutes with shaking. Then the solvent was removed, and gel pieces were allowed to air dry for 20 min. To the dry gel pieces, 10 mM TCEP and 40mM chloroacetic acid were added and incubated at 70ºC for 5 min. The gel pieces were washed again with 50% acetonitrile and 50% 50mM ammonium bicarbonate for 15 min with shaking. The gel pieces were then rehydrated in 50 mM ammonium bicarbonate and 1ug Chymotrypsin/GluC (1:50) was added and incubated for 1 hour at room temperature, then 50mM HEPES pH 8 solution was added to cover the pieces. The samples were allowed to incubate overnight at 37ºC. Peptides were extracted from gel pieces with 25% acetonitrile and 50mM ammonium bicarbonate pH 8, then again with 100% acetonitrile for 5 min with shaking. Samples were filtered through a 0.22 µm PVDFspin column (Millipore) Peptides were dried in a speedvac apparatus to 30-60 µl total volume, then samples were acidified with 2 µl formic acid (neat).

In-gel Chymotrypsin/GluC digested peptides were analyzed by online capillary nano LC-MS/MS using a 25 cm reverse phase column and a 10 cm precolumn fabricated in-house (75 µm inner diameter, packed with ReproSil-Gold C18-1.9 μm resin (Dr. Maisch GmbH)) that was equipped with a laser-pulled nanoelectrospray emitter tip. The precolumn used 3.0 µm packing (Dr. Maisch GmbH)). Peptides were eluted at a flow rate of 300 nL/min using a linear gradient of 2–40% buffer B in 140 min (buffer A: 0.05% formic acid and 5% acetonitrile in water; buffer B: 0.05% formic acid and 95% acetonitrile in water) in an Thermo Fisher Easy-nLC1200 nanoLC system. Peptides were ionized using a FLEX ion source (Thermo Fisher) using electrospray ionization into a Fusion Lumos Tribrid Orbitrap Mass Spectrometer (Thermo Fisher Scientific). Instrument method parameters were as follows: MS1 resolution, 120,000 at 200 *m/z*; scan range, 350−1600 *m/z*, orbitrap mode acquisition. The top 20 most-abundant ions were subjected to collision-induced dissociation with a normalized collision energy of 35%, activation q 0.25, and precursor isolation width 2 *m/z*. Dynamic exclusion was enabled with a repeat count of 1, a repeat duration of 30 seconds, and an exclusion duration of 20 seconds. RAW files were analyzed using PEAKS (Bioinformatics Solution Inc) with the following parameters: semi-specific cleavage specificity at the C-terminal site of R and K, allowing for 5 missed cleavages, precursor mass tolerance of 15 ppm, and fragment ion mass tolerance of 0.5 Daltons. Methionine oxidation was set as variable modifications and cysteine carbamidomethylation was set as a fixed modification. Peptide hits were filtered using a 5% FDR. Proteins with at least 2 unique peptides were filtered with a 5% FDR. Label free quantitation (LFQ) was performed using the PEAKS quantitation module and default parameters with the following exceptions: Top 2 peptides for each protein with a min of 10XE4 abundance was used and the TIC was used for all normalization including technical replicates. Peptide mass spectrometry data were visualized using PrIntMap-R^63^. See Tables S3-5 for processed data containing reads from chymotrypsin and GluC digested peptides used to generate Fig. 2E.

### Protease-HamAB nuclease plasmid clearance assay

Plasmid interference assays were conducted in isothermal amplification buffer. Plasmids were diluted to 4 ng/μl. Nicked and cut plasmid controls were generated by treatment with Nt.BspQI (New England Biolabs) and BamHI-HF (New England Biolabs), respectively. Reactions were started with addition of Protease-HamAB at 5 nM and HamM at 10 nM, and incubated at 30°C for 1 min, 5 min, or 30 min. Reactions were quenched with addition of proteinase K (New England Biolabs) and 10x Bluejuice. Reactions were imaged on 0.5% TAE agarose gels. Gels were stained with SYBR Safe and imaged on a ChemiDoc MP (BioRad).

### Protease-HamAB nuclease RNAse activity assay

RNA cleavage assays were performed in isothermal amplification buffer or MBP/SEC buffer supplemented with 1mM ZnCl_2_. 0.5 µM of protease-HamAB and 0.5 µM of HamM were incubated with 0.5 µM dsRNA or 0.5 µM ssRNA for 30 minutes at 30°C. Reactions were stopped by addition of Proteinase K (New England Biolabs) and 10x Bluejuice. Reactions were run on a 2% agarose gel stained with SYBR Gold (Thermo Fisher) and imaged on a ChemiDoc MP (BioRad). For a summary of oligonucleotide substrates, see Table S6.

### Malachite green ATPase assays

Orthophosphate liberation was determined with a Malachite Green Phosphate Assay kit (BioAssay Systems) according to the manufacturer protocol. All reactions were conducted in isothermal amplification buffer. Substrates tested included two different ssDNAs (ssDNA 1 and ssDNA 2), blunt-ended dsDNA, dsDNA with 3′ overhang, dsDNA with 5′ overhang, and plasmid DNA. DNA substrates were diluted to 100 nM, or 4 ng/µl for plasmid reactions, in a total reaction volume of 40 µl in clear bottom, flat, black 96-well assay plates (Corning Costar). Reactions were started with addition of *Pfl*HamAB to 40 nM and ATP to 1 mM, followed by incubation at 30°C for 30 minutes. Additional controls with buffer only, ATP only, HamAB only, and no DNA substrates were performed. Reactions were quenched with the addition of activated malachite green reagent. The absorbance values of wells were measured after 20 min of color development at ambient temperature with a Biotek plate reader at 620 nm. Orthophosphate liberation was interpolated against a standard curve with known concentrations of free phosphate. For a summary of oligonucleotide substrates, see Table S6.

### Nuclease and caspase structural analyses

Nuclease structures from *Pfl*HamM, *Epi*Avs1a and *Kae*CanC were predicted using the AlphaFold3 web server^23^. Predicted structures were compared to the AlphaFold databank (AFDB) using Foldseek^64^. A confident hit, (uniprot: P39695) protein 3 from the *B. subtilis* ComE operon, was aligned and visualized with matchmaker in ChimeraX^58^. Sequence logos of insertion regions containing the putative protease cleavage sites as determined by mass spectrometry (see above) were generated by performing BLAST searches, then aligning 100 hit sequences with ClustalOmega^50^ and visualizing regions of interest with the built-in sequence logo function of Geneious Prime. To generate a structural comparison of *Kae*CanA, Dali was used to search the protein data bank for remote homology to experimentally determine structures, then superpositions were regenerated and visualized in ChimeraX using matchmaker.

