## Supplementary References and Figures for "Recurrent acquisition of nuclease-protease pairs in antiviral immunity"

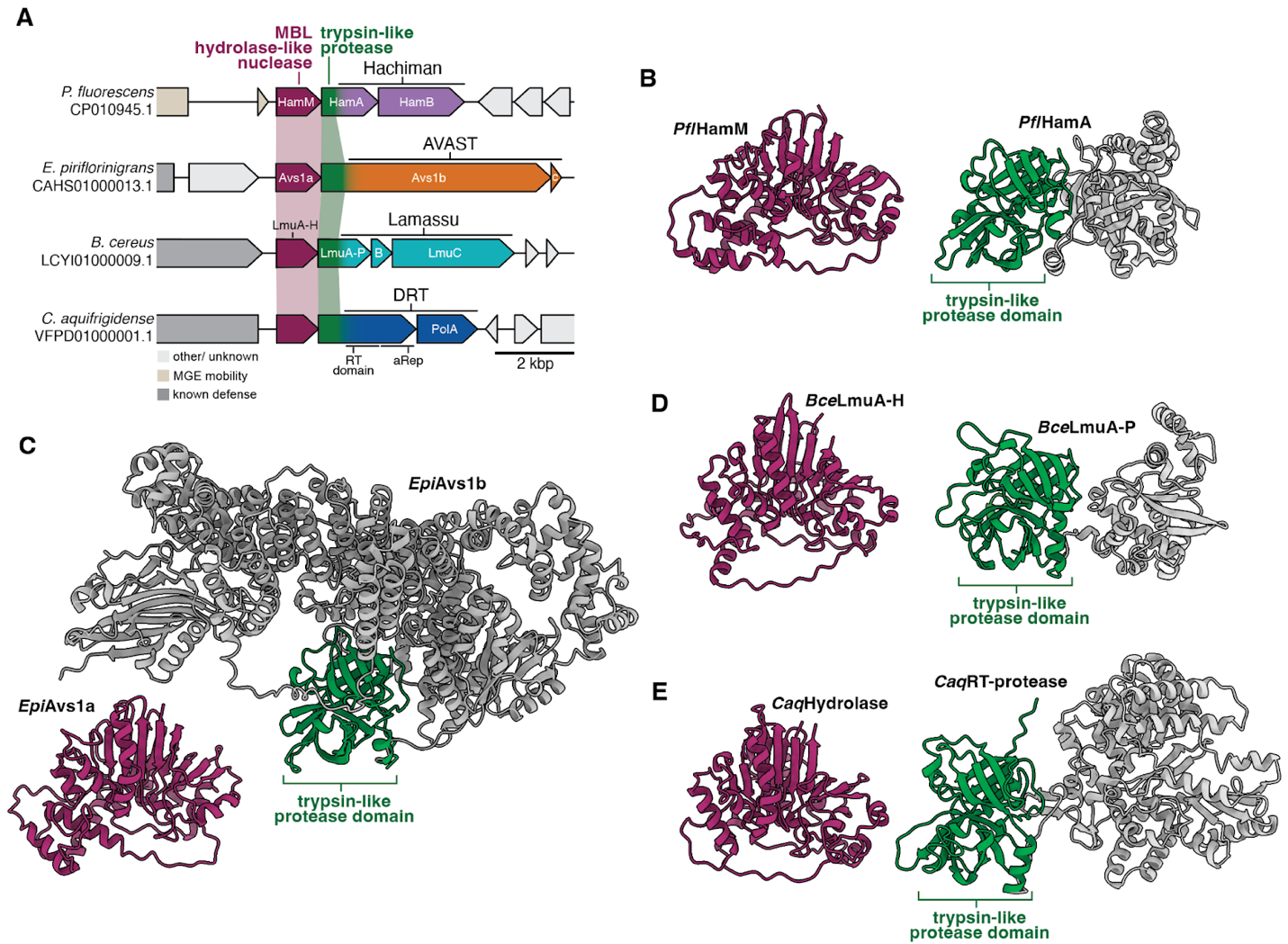

**Fig. S1. Structural comparison of nuclease-protease pairs in known defense systems.** (**A**) Loci illustrating the genetic architecture of multiple defense systems encoding an N-terminal MBL hydrolase-like nucleases and partnering tyrpsin like protease, as in Fig. 1A. Annotations for gene context and gene names are shown. (**B**-**E**) AlphaFold 3 predicted protein structures for nuclease and protease-fusion genes in Hachiman, AVAST, Lamassu, and DRT (UG9) systems. In each case, protein structures were predicted using loci from (A). Predicted structural snapshots are aligned to show homology.

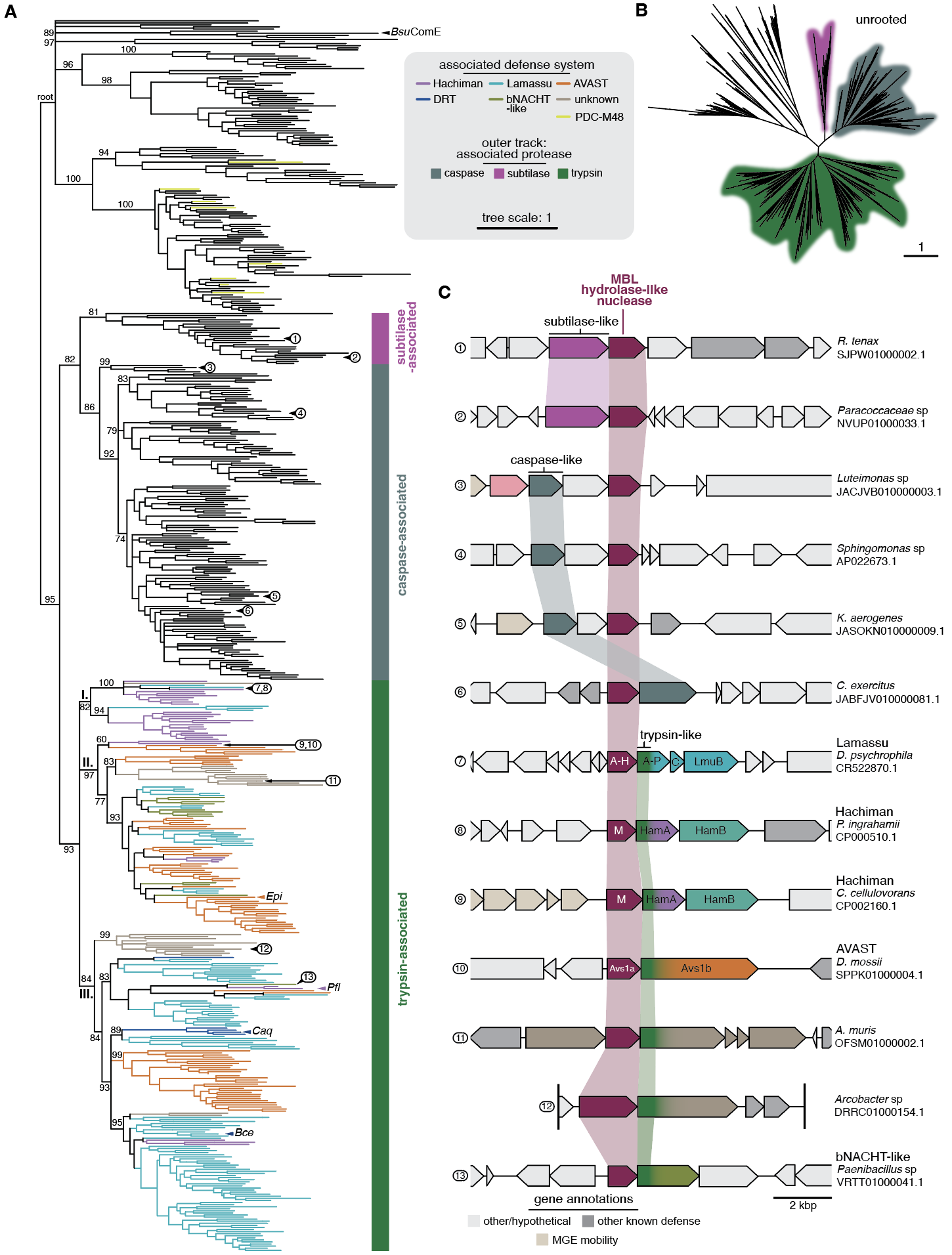

**Fig. S2. Modular evolution of defense-associated MBL hydrolase-like nucleases.** (**A**) Phylogenetic tree of diverse MBL hydrolase-like nuclease sequences in bacteria. The phylogenetic tree is rooted at a large clade containing MBL hydrolase-like nucleases without known protease associations which encode ComE nuclease domains. Tree leaves are colored according to inclusion of the predicted nuclease in known defense systems as determined by PADLOC search of a 20 kbp genomic context. The outer track indicates association with nearby protease families. Three major operonic protease families - subtilase-, caspase-/CHAT- and trypsin-family proteases were detected. Branches with support values < 60 are deleted, resulting in polytomy. Select bootstrap support values are shown. The clade containing trypsin-associated nucleases is shown as a subtree in Fig. 1B. A subtree of protease-associated nucleases is shown with clades collapsed in Fig. 4A. (**B**) Unrooted form of phylogenetic tree from (A). Colored ranges indicate associated proteases as in (A). (**C**) Genomic loci of nucleases from the phylogeny in (A). Select loci from each major protease-associated clade are labelled in A. Genes, proteases and detected defense systems are annotated as in (A) and according to the key below.

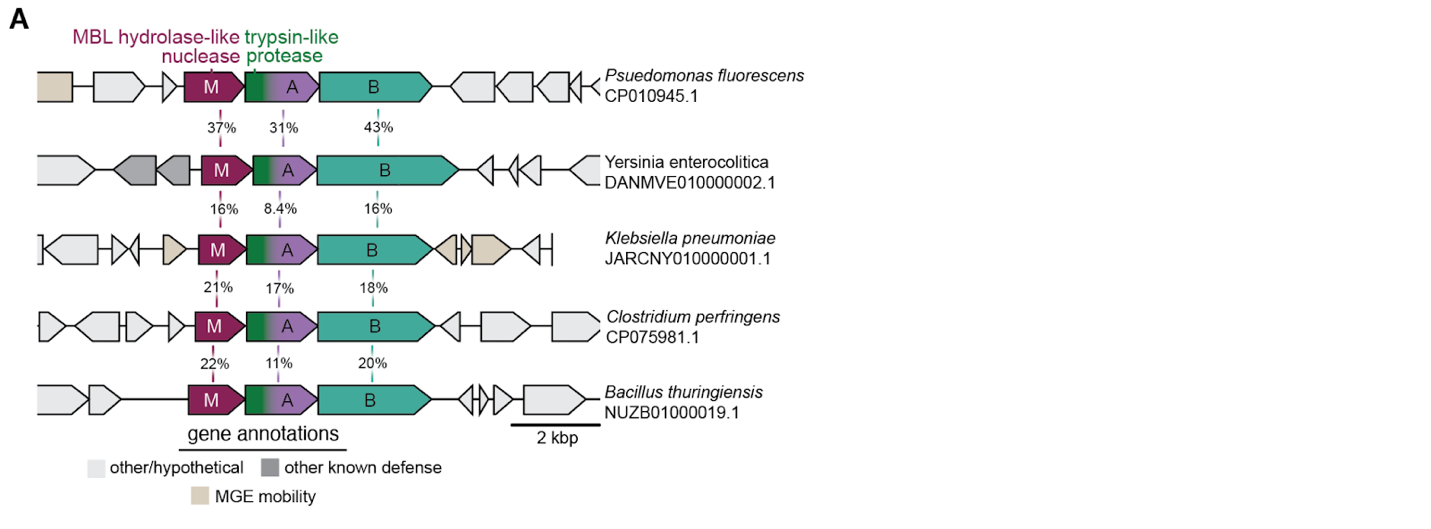

**Fig. S3. Nuclease-protease Hachiman systems are highly diverse.** (**A**) Genomic loci encoding nuclease-protease Hachiman systems from different clades of the nuclease phylogeny in fig. S2A. Percent identity between different genes are shown between loci. *P. fluorescens* and *Y. enterocolitica* sequences originate from the same clade.

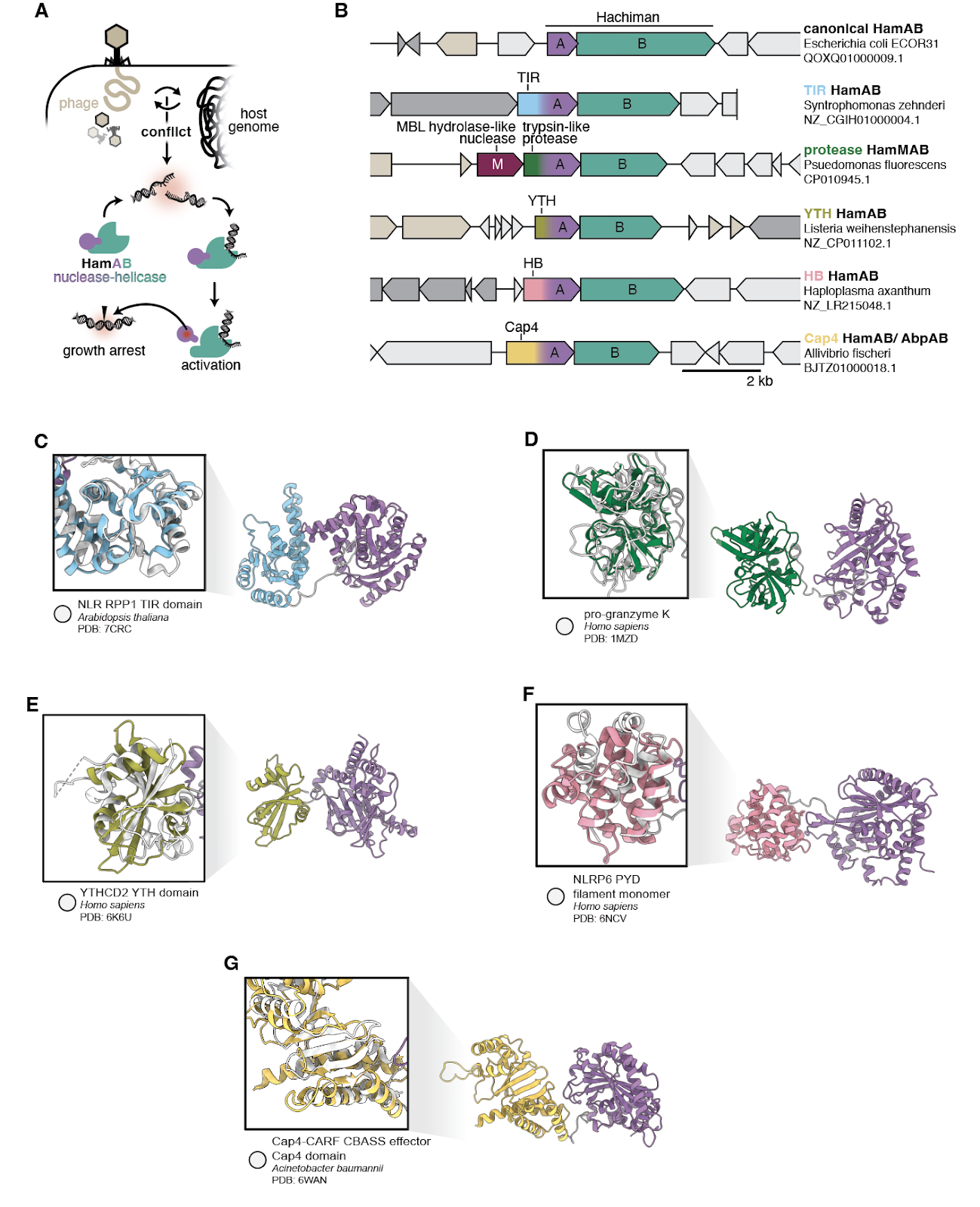

**Fig. S4. Hachiman HamA acquired multiple additional domains.** (**A**) Model for Hachiman antiphage defense. (**B**) Loci illustrating the genetic architecture of canonical and fusion variant Hachiman systems. The five fusion-HamA systems identified are: Toll/interleukin-1 receptor (TIR) domain-HamA, trypsin protease-HamA with upstream metallo‐β‐lactamase fold hydrolase-like nuclease (HamM), YT521-B homology (YTH) domain-HamA, helical bundle (HB) domain-HamA, and CBASS activated protein 4 (Cap4) domain-HamA or AbpA. (**C**-**G**) AlphaFold structural predictions of domain-HamA fusions shown in (B). Inset boxes show structural superpositions with DALI-predicted homologous experimental structures, which are shown in white and are indicated below in each case.

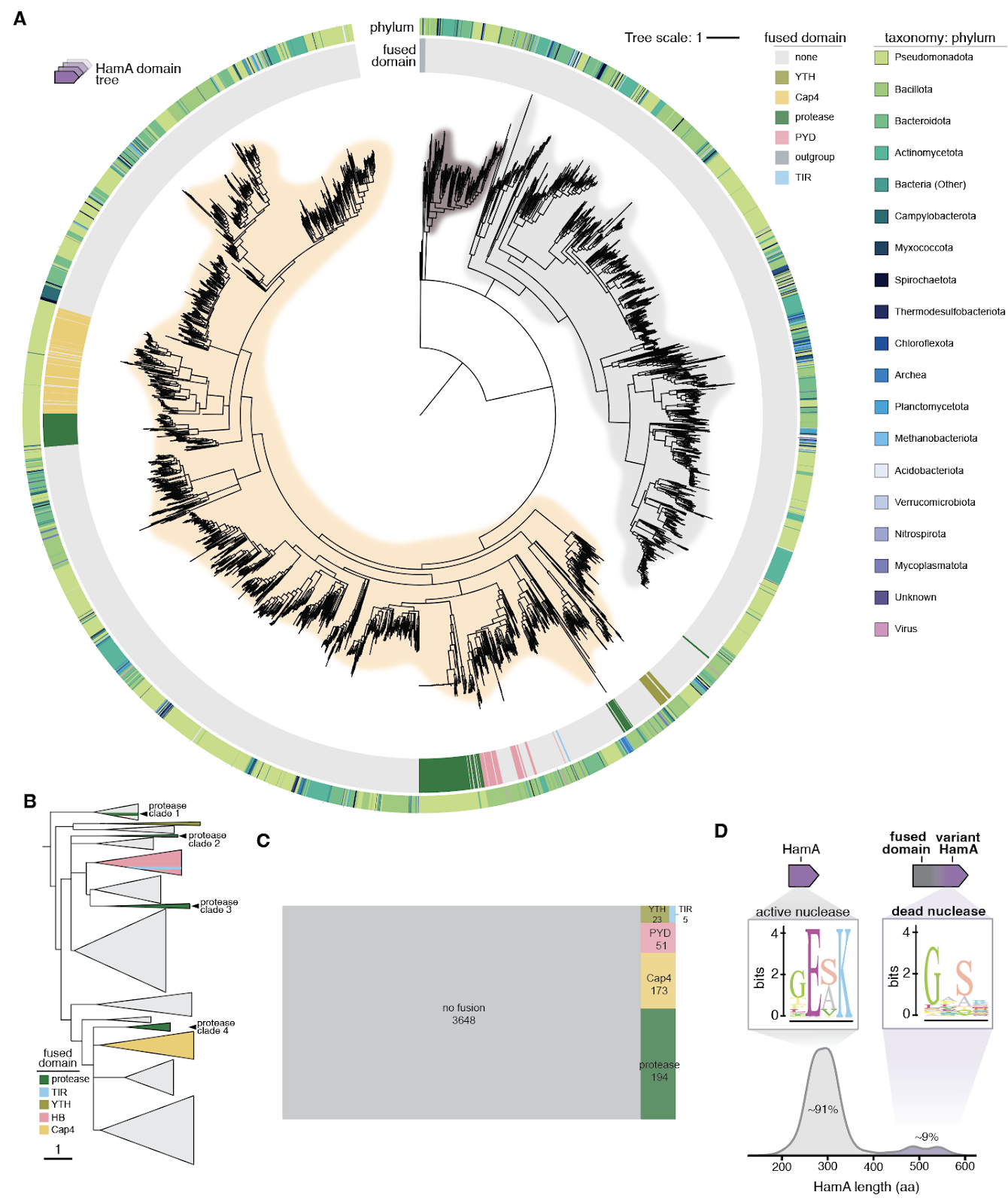

**Fig. S5. HamA domain phylogeny and loss of nuclease activity.** (**A**) Phylogenetic tree of 4,094 HamA domain sequences. The tree is rooted at a clade containing 10 homologs of the *P. aquatile* type IIS restriction endonuclease domain. The inner track indicates presence of a fusion domain, but these were excluded from phylogenetic analysis to ascertain the phylogeny exclusively of the HamA domain. The outer track indicates associated phyla. Branches with support values < 60 are deleted, resulting in polytomy. (**B**) Cartoon of the fusion containing clade in (A), with clades collapsed and color-coded to indicate fusion content. (**C**) Fraction of HamA sequences encoding fusions. (**D**) Sequence logos of the nuclease active site for canonical versus domain fusion HamA variants are shown above a density plot showing the distribution of HamA protein lengths across HamA diversity. HamA variants were manually curated from all HamA domain sequences from panel (A) and aligned.

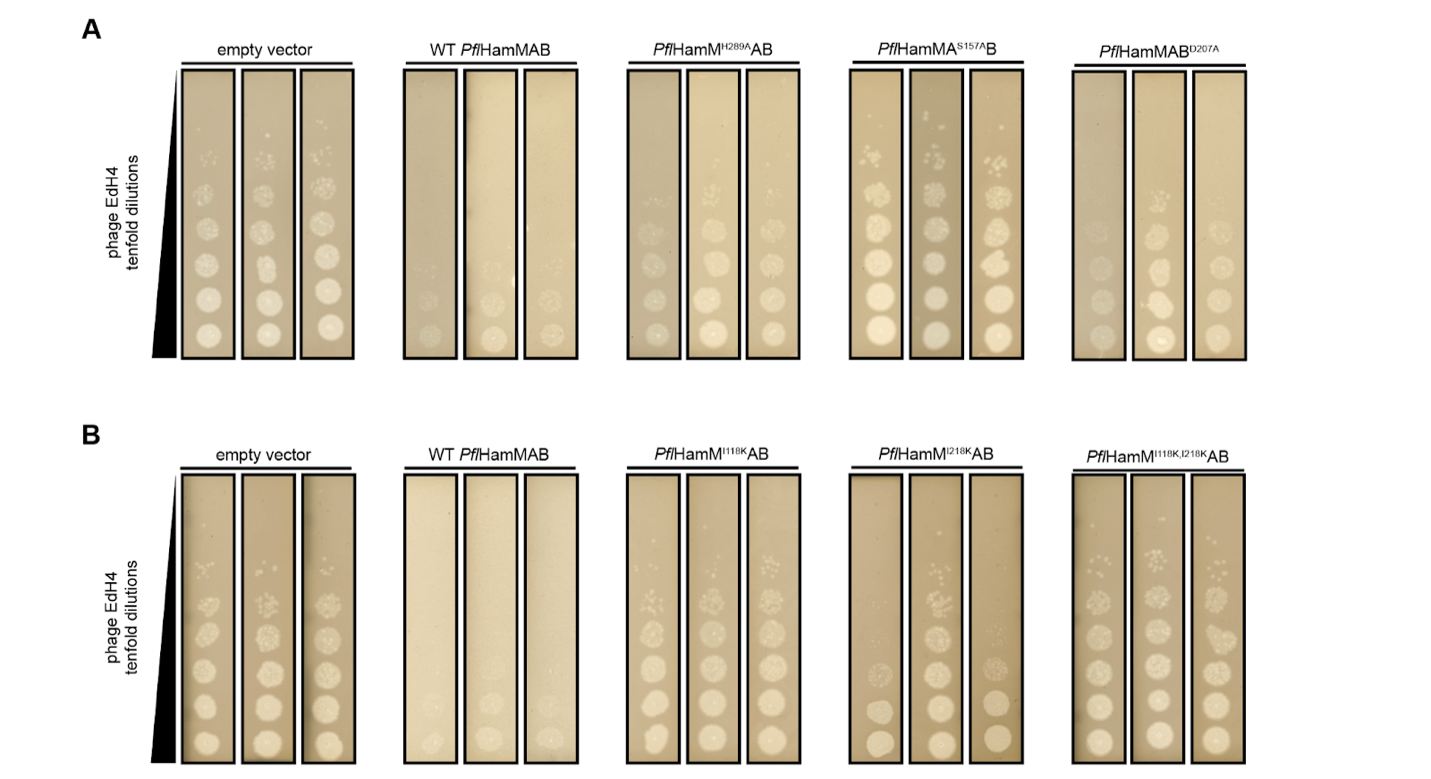

**Fig. S6. Plaque assays of *P. fluorescens* HamMAB and mutants.** (**A**) Images of triplicate plaque assays where *E. coli* expressing *Pfl*HamMAB and catalytic mutants were challenged with tenfold dilutions of phage EdH4. (**B**) Images of triplicate plaque assays where *E. coli* expressing *Pfl*HamMAB and cut site mutants of HamM were challenged with tenfold dilutions of phage EdH4.

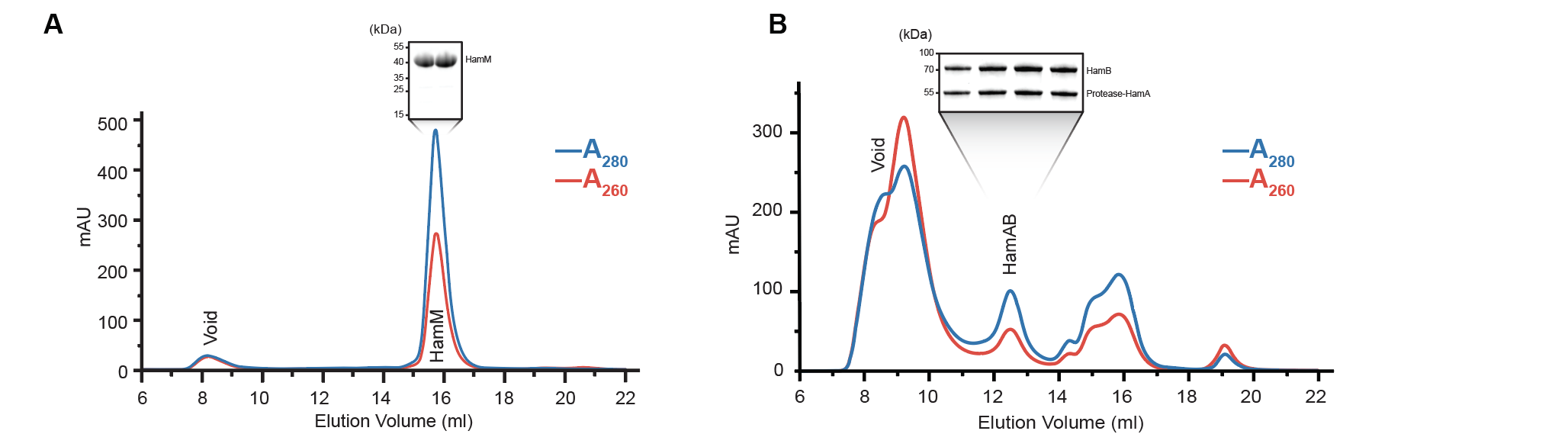

**Fig. S7. Purification of HamM and HamAB. (A)** Size exclusion chromatography trace of HamM after TEV protease treatment and removal of MBP by reverse immobilized metal affinity purification (IMAC). Corresponding peaks of elution fractions are run on a Coomassie PAGE gel. **(B)** Size exclusion chromatography trace of HamM after TEV protease treatment and removal of MBP by reverse IMAC. Corresponding peaks of elution fractions are run on a Coomassie PAGE gel.

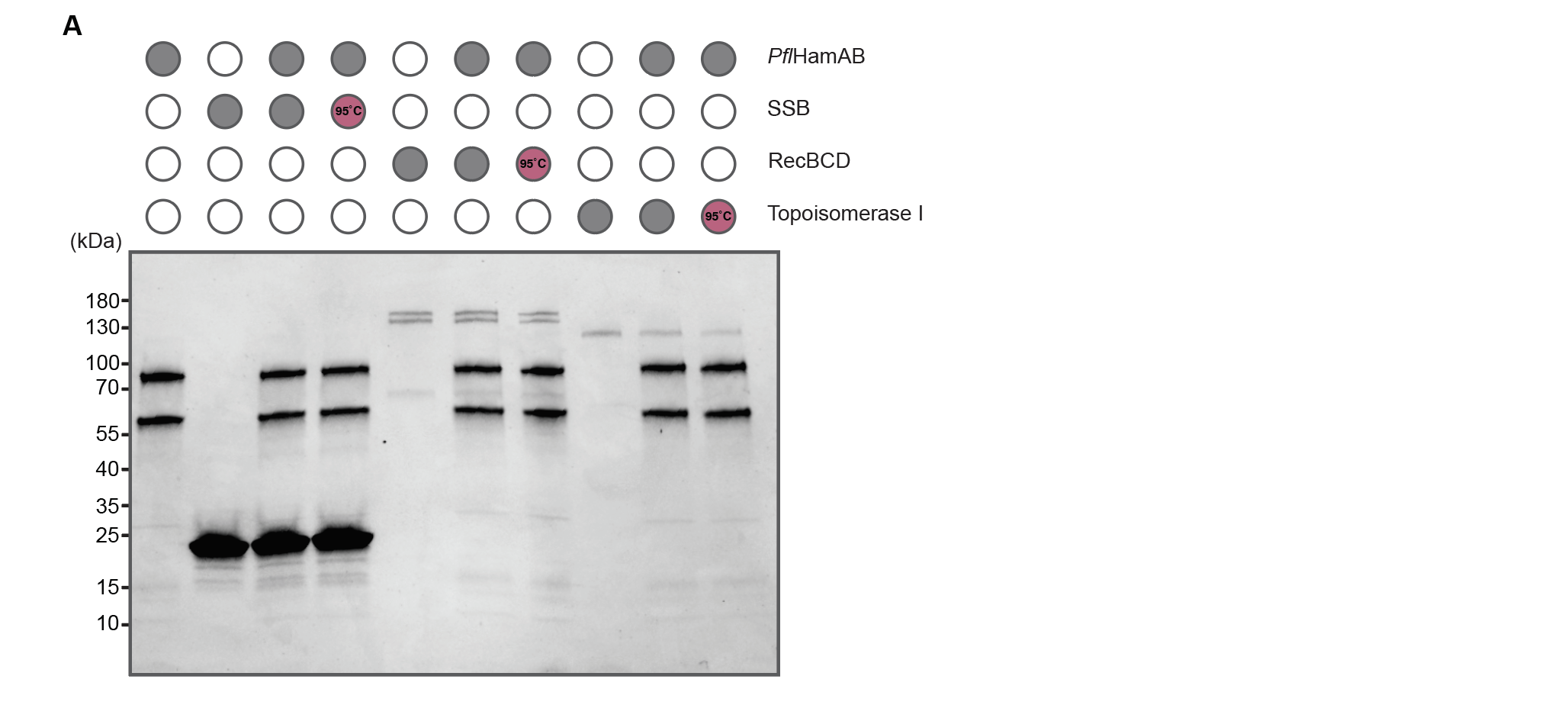

**Fig. S8. *In vitro* incubation of *Pfl*HamAB with Single-Stranded DNA Binding Protein (SSB), RecBCD, and Topoisomerase I. (A)** *In vitro* reactions of HamAB with SSB, RecBCD, and Topoisomerase I visualized on a Coomassie PAGE gel. In lanes 4, 7, and 10 (from left to right), protein substrates were pre-boiled before addition of HamAB.

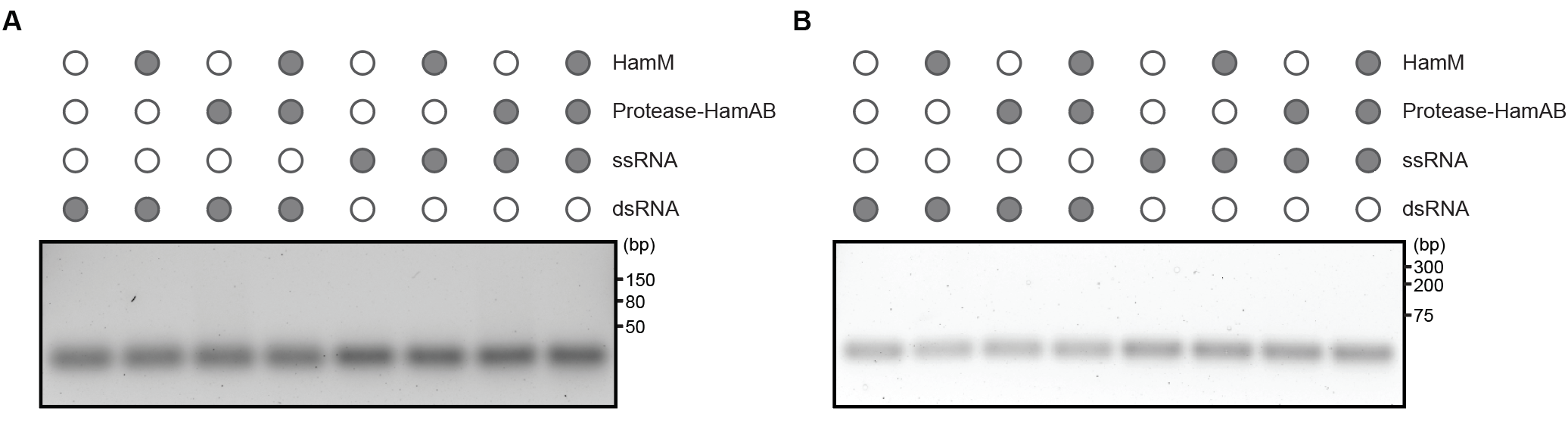

**Fig. S9. *In vitro* RNA cleavage assay of *Pfl*HamMAB. (A)** *In vitro* reactions of HamM and HamAB against dsRNA and ssRNA in isothermal amplification buffer, visualized on a 2% agarose gel stained with SYBR Gold. **(B)** *In vitro* reactions of HamM and HamAB against dsRNA and ssRNA in MBP/SEC Buffer supplemented with 1mM ZnCl_2_, visualized on a 2% agarose gel stained with SYBR Gold. For a summary of oligonucleotide substrates, see Table S6.

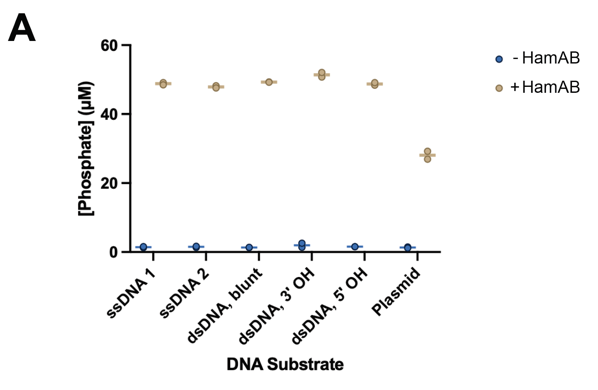

**Fig. S10. *In vitro* ATPase assay of *Pfl*HamAB. (A)** Malachite green ATPase assays of HamAB with ATP against a panel of DNA substrates. Individual data points of two independent biological replicates are shown. For a summary of oligonucleotide substrates, see Table S6.

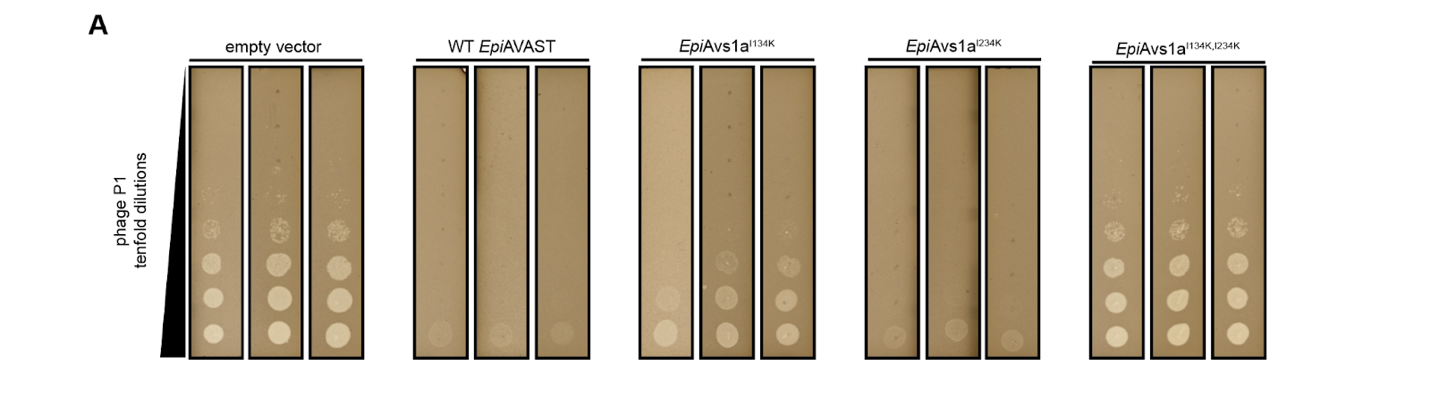

**Fig. S11. Plaque assays of *E. piriflorinigrans* AVAST and Avs1b mutants.** (**A**) Images of triplicate plaque assays where *E. coli* expressing *Epi*AVAST and catalytic mutants of Avs1a were challenged with tenfold dilutions of phage P1.

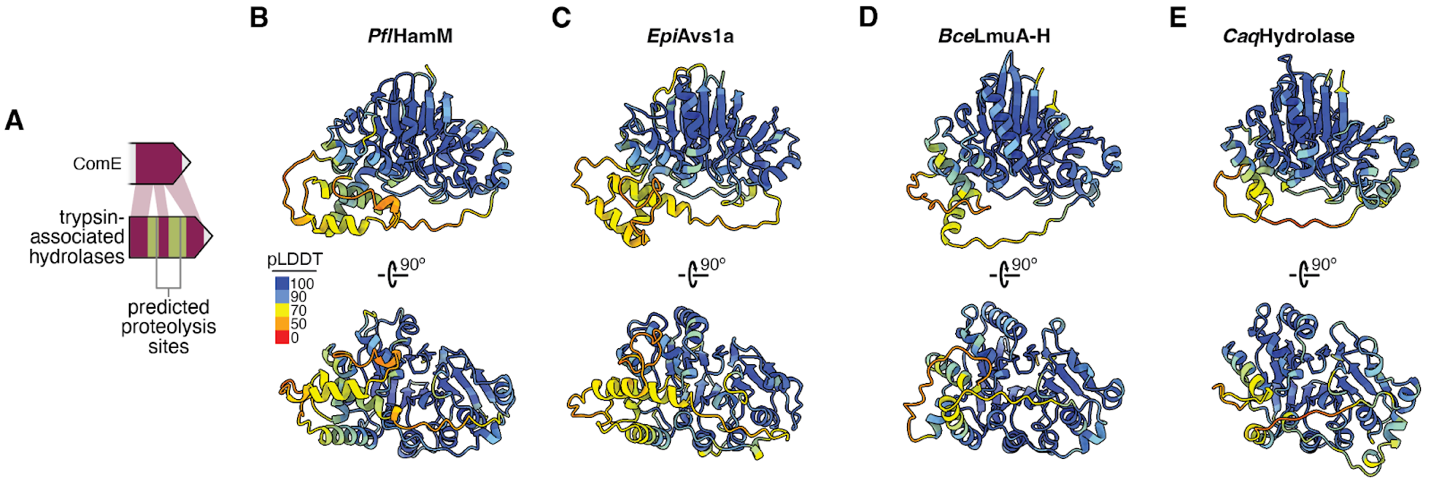

**Fig. S12. Structural comparisons of trypsin-associated nucleases.** (**A**) Model for the emergence of autoinhibitory ‘gating’ insertions (green) in trypsin-associated nucleases in antiphage systems. (**B**-**E**) AlphaFold3 predicted protein structures for nuclease genes in Hachiman, AVAST, Lamassu and DRT (UG9) systems, as in fig. S1 but with an additional view shown and colored by pLDDT. In each case, protein structures were predicted using loci from Fig. 1a. Predicted structural snapshots are aligned to show homology.

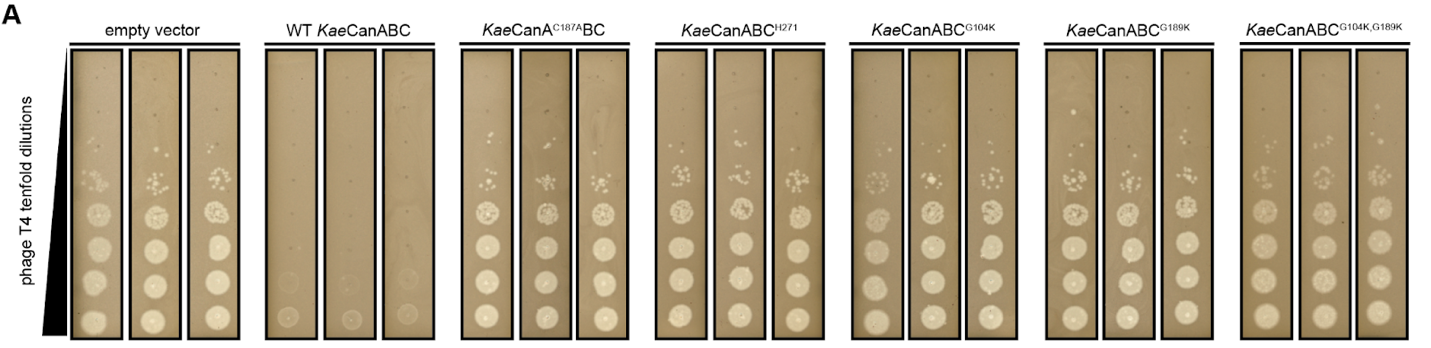

**Fig. S13. Plaque assays of *K. aerogenes* CanABC and mutants.** (**A**) Images of triplicate plaque assays where *E. coli* expressing *Kae*CanABC and various catalytic and cut site mutants were challenged with tenfold dilutions of phage T4.

**Table S1. (separate file)**

Cloned sequences used in this study.

**Table S2. (separate file)**

Summary of plasmids used in this study.

**Table S3. (separate file)**

Mass spetrometry results for HamM fragment 1 (high molecular weight).

**Table S4. (separate file)**

Mass spetrometry results for HamM fragment 2 (medium molecular weight).

**Table S5. (separate file)**

Mass spetrometry results for HamM fragment 3 (low molecular weight).

**Table S6. (separate file)**

Oligonucleotide constructs used in biochemical assays.
